# Salinity plays a limited role in determining rates of size evolution in fishes globally across multiple scales

**DOI:** 10.1101/2023.09.05.556281

**Authors:** John T. Clarke, Robert B. Davis

**Affiliations:** GeoBio-Center, Ludwig-Maximilians-Universität München, Munich, Germany; Department of Earth and Environmental Sciences, Palaeontology & Geobiology, Ludwig-Maximilians-Universität München, Munich, Germany; Institute of Ecology and Earth Sciences, Department of Zoology, University of Tartu, Tartu, Estonia; Department of Ecology and Biogeography, Faculty of Biological and Veterinary Sciences, Nicolaus Copernicus University, Toruń, Poland

## Abstract

Substantial progress has been made to map biodiversity and its drivers across the planet at multiple scales, yet studies that quantify the evolutionary processes that underpin this biodiversity, and test their drivers at multiple scales, are comparatively rare. Studying most fish species, we quantify rates of body size evolution to test the role of fundamental salinity habitats in shaping rates of evolution at multiple scales. We also determine how four additional factors shape evolutionary rates.

In up to 1710 comparisons studying over 27,000 ray-finned fish species, we compare rates of body size evolution between five salinity habits using 12 metrics. The comparisons span a molecular tree, supertrees, and ten scales of observation to test for robust patterns and reveal how patterns change with scale. Then, three approaches assess the role of three non-salinity factors on rates, and an alternative habitat scheme tests if lakes influence evolutionary rates.

Rates of size evolution rarely differ consistently between salinity habitats; rate patterns are highly clade- and scale-dependent. One exception is freshwater-brackish fishes, which possess among the highest size rates of any salinity, and show higher rates than euryhaline fishes in most groupings studied at most scales, and verses marine, freshwater, and marine-brackish habitats at specific scales. Additionally, species richness had the greatest potential to predict phenotypic rates, followed by branch duration and absolute values of body size. Lacustrine environments were consistently associated with high rates of size evolution.

We reveal the rate patterns that underpin global body size diversity for fishes, identifying factors that play a limited role in shaping rates of size evolution, such as salinity, and those such as species richness, age, and lake environments that consistently shape evolutionary rates across half of vertebrate diversity.

## INTRODUCTION

Understanding the mechanisms that explain biodiversity on the planet, including how and why these mechanisms vary with scale, is essential to both evolutionary biology and macroecology. Substantial progress has been made to recognise (Blackburn and Gaston, 2002) and document (Clarke, 2021; Finlay et al., 2006; Foody, 2004; McClain and Nekola, 2008; Rahbek, 2005) patterns and drivers of present-day biodiversity at multiple scales. By contrast, similarly broad efforts to understand what factors drive differences in the evolutionary processes that underlie this biodiversity (e.g. rates of phenotypic evolution), and how these vary with scale, are comparably rare, despite their ability to deliver significant insights. For example, the positive relationship between speciation rates and rates of size evolution at the global scale is frequently weak or absent at lower taxonomic scales (Cooney and Thomas, 2020), and different functional traits (e.g. body size relative to jaw related feeding traits) are more or less likely to differ between taxa depending upon the scale (e.g. genus, family or higher-level scales) at which the analysis is performed (Slater and Friscia, 2019). Containing over half of vertebrate diversity (34,000+ species) and six orders of magnitude in body size variation, fishes possess huge potential to deliver highly generalisable results regarding how ecological conditions shape phenotypic evolution and how this varies with scale. Furthermore, fishes are uniquely well positioned to test drivers of evolutionary mechanisms in aquatic settings, given they occupy nearly every aquatic environment.

One of the most fundamental factors thought to shape biodiversity and underlying evolutionary processes in the aquatic realm is the divide between marine and freshwater environments, owing to the rarity of organisms that can successfully cross this divide (Lee and Bell, 1999). Fishes represent an exception to this general rule, with many groups having regularly transitioned between these environments at both small (e.g. within families) and large (e.g. between orders) phylogenetic scales. This repeatability at multiple scales provides an ideal ‘natural experiment’ in which to elucidate the role these habitats play on rates of size evolution at multiple scales of observation. While progress has been made testing the role of salinity habitat on speciation rates at broad scales (Miller, 2021; Rabosky, 2020), analogous broad-scale approaches for rates of size evolution are lacking. A number of valuable case studies do permit comparisons between marine and freshwater environments (Betancur-R. et al., 2012; Davis et al., 2014; Guinot and Cavin, 2018; Santini et al., 2013; Thacker, 2014), but their differing methods, phylogenetic scales, and results make it difficult to draw overall conclusions. There is, however, compelling evidence that absolute body size does vary systematically between salinity environments, especially when the question is broadened from a binary marine vs. freshwater comparison to consider marine-brackish, freshwater-brackish, and euryhaline salinity lifestyles (Clarke, 2021). Specifically, actinopterygian fishes show a three-tiered size distribution according to salinity habitat, with small freshwater taxa, medium-sized freshwater-brackish and marine taxa, and large euryhaline and marine-brackish taxa. These systematic differences in absolute size between habitats beg the question as to whether these habitats are underpinned by different underlying rate dynamics.

Our intention, with a ∼10,000 species molecular tree, and one hundred ∼26,000 supertree datasets of body size and salinity habitat for actinopterygian fishes, is to test whether rates of size evolution differ between salinity habitats when examined across ten phylogenetic scales. This allows us to overcome ascertainment bias and reveal the overall architecture of rate patterns, showing how these rates differ within and between scales, and determining whether results for the whole dataset are underpinned by consistent outcomes at lower scales.

We used both a phylogeny that only includes taxa with molecular data, and supertrees that add the rest of fish diversity using taxonomic data, to help us to evaluate whether a recently demonstrated size bias in the molecular taxon sample towards larger absolute body sizes (Clarke, 2021) has a discernible impact upon rates of size evolution. In addition, we use up to 1056 comparisons and the full dataset to evaluate the role of three additional variables on rates of size evolution. Finally, we examine an alternative aquatic habitat scheme to explore how lakes and rivers impact phenotypic evolution.

## MATERIALS AND METHODS

### Data, phylogenetic scales for comparisons, and molecular versus supertree analyses

We conduct our analyses using both one 11,638-tip molecular tree of actinopterygians and the 100 supertrees constructed from this molecular backbone (Rabosky et al., 2018), referred to as the 11k tree and 31k trees herein. All species for which salinity data were available in FishBase (Froese and Pauly, 2000) were then assigned to the six categories: i. exclusively marine; ii. exclusively freshwater; iii. exclusively brackish; iv. marine-brackish, v. freshwater-brackish; vi. euryhaline. Exclusively brackish taxa were not subject to detailed pairwise comparisons as they possess extremely few species. Two datasets were constructed – a 10,906-species dataset with size and salinity data matching taxa in the 11k tree, and a 27,226-species dataset with size and salinity data matching taxa in the 31k tree). Analysis files are available on Dryad. Analyses were run on both 11k- and 31k-tree matched datasets. Both dataset types have potential biases, as body sizes are biased towards larger size in molecularly sampled taxa (Clarke, 2021), while supertrees may also alter phenotypic rate results (Rabosky, 2015); thus, it is important to consider both.

We design the study to overcome ascertainment bias (i.e. where clades that are more varied in the trait of interest, here salinity habitat, are more likely to be studied). This is important, as “hotspot” clades for salinity variation in fishes (i.e. clades that explore many salinity habitats) were more likely to show unusual patterns of absolute size differences between salinity habitats relative to clades we would not consider hotspots (SI text of Clarke, 2021). To avoid this, we test for rate differences within all definable groups at ten different phylogenetic scales. This also allows us understand how rates propagate at different scales, which has been demonstrated to be highly important for macroevolutionary results (Clarke, 2021; Rabosky, 2020), especially where specific traits (e.g. feeding traits) are more likely to change at specific scales (Slater and Friscia, 2019). Phylogenetic scales 1 through to 7 test patterns at increasingly broad phylogenetic scales, first testing all fish families (1), then all orders (2), then successively larger groupings on the phylogeny termed Tax3, Tax4, Tax5 and Tax6 (where the phylogeny is split into 13, 9, 5, and 3 sections, respectively), followed by the full dataset (7). The final three scales have a different goal, studying only those groupings on the phylogeny considered “evolutionary hotspots” for salinity as a demonstration of ascertainment bias (8), then expanded versions of these hotspots to consider whether species surrounding the hotspot had a mediating effect (9), and order-level groups with hotspot orders removed (10) to consider if more typical clades showed different dynamics to hotspots.

### Testing for differences in size rates between salinity habitats

For all unique habitat+group comparisons made in this study (1056 and 1710 for the 11k-tree and 31k-tree datasets, respectively), 12 metrics are used to compare rates of size evolution. The first five of these metrics pertain to those obtained from non-log-transformed BAMM tip-rate data. BAMM uses a Bayesian framework to estimate shifts in rates of trait evolution across a phylogeny within a Brownian motion framework (Rabosky et al., 2014). Convergence was achieved for the 11k-tree run for 1 billion generations, sampling every 380,000 generations, with the first 15% of samples as burn-in). To achieve convergence on the 31k supertree, it was split into six segments, and each was run for 1.5 billion generations, sampling every 375,000 generations. The segments were then merged using the “add_subclade_to_backbone_BAMM” function (Igea et al., 2017). Priors were informed by the SetBAMMPRIOR function. Convergence was checked with R package coda. The five metrics derived from this data are: i. mean BAMM tip-rates, and the use of these tip-rates to compute the following four metrics, ii. a Wilcoxon test, iii. phylogenetic averages with PGLS ANOVA, R package RRPP (Collyer and Adams, 2018), and iv. the PGLS ANOVA’s significance test, and v. simulation phylogenetic ANOVA significance tests, R package phytools (Revell, 2012). The analogous five metrics (vi–x) were then obtained for log10 mean BAMM tip-rates.

Metric xi. uses the compare.rates function in the R package geomorph (Adams and Otárola-Castillo, 2013) once for every comparison in the 11k-tree dataset, and then over the 100 topologies of the 31k-tree datasets (except for the largest phylogenetic scales, which are run once due to computational demands). Metric xii. is the significance test performed by STRAPP in BAMMtools (Rabosky and Huang, 2016). Together, these thousands of results allow the reader to examine any specific metric or clade comparison of interest (Appendices 1–4) and allow us to test for robust patterns that hold across datasets and scales. BAMM runs together with other rate comparisons took several months to compute.

### Testing alternative drivers of rate variation

Beyond salinity habitat, we tested three alternative predictors of rate (species richness, tip-branch duration, absolute size) with three approaches (A–C in Table 1, details below), allowing us to examine both variables assigned to groups (e.g. species richness) and those assigned to species (e.g. body size). Considering results from all three approaches, associations are then ranked by their degree of support (Table 1). Approaches A and B require us to define groupings of taxa. For this, we used the same 125 unique habitat comparisons at the order level which were previously used to test for differences in rate between the six main salinity habitats in the 11k-datasets (i.e. every comparison in the Ord. column totals to 125, e.g. Appendix 1).

**TABLE 1:** Three hypothesised associations between rates of body size evolution and predictor variables, ranked from 1 (highest support) to 3 (least support) after considering results from three methods designed to assess the strength of these associations.

| Approach A: Alignment of discrete outcomes |  | Approach B: Continuous relationship |  | Approach C:<br>Continuous relationship |
| --- | --- | --- | --- | --- |
| Performed using 125 habitat comparisons at the order level<br>Appendix 5 shows where each hypothesised association, and<br>Appendix 6 shows % summaries of the order level comparisons. |  | Assessed with regressions for 10 sets of pairwise<br>habitat comparisons at order level.<br>See Appendices 7, 8 and 9 to view each regression,<br>and Appendix 10 for statistical summaries. |  | Assessed with STRAPP<br>using species-specific<br>values. |
| 1. Testing the hypothesised positive association between **phenotypic rates** and **species richness**.<br>May expect, on average, that a group with more species has experienced higher speciation rates, and thus, would also possess higher phenotypic rates<br>(given their previously established correlation in fishes). **Positive association not supported. Instead, a negative association has <u>moderate support</u>.** |  |  |  |  |
| Only 48/125 (38.4%) of comparisons support a positive association<br>Pos. association is particularly weak when comparing the specific<br>habitats, where only 2/10 (20%) of Freshwater vs. Marine-brackish<br>comparisons, and 3/15 (20) Euryhaline vs. Marine-brackish support a<br>positive association. | | 2/10 regressions support a positive association.<br>The 2 regressions have a mean $R^2 = 0.164$ and mean<br>$p = 0.228$<br>**Instead, 8/10 regressions support negative**<br>**association.** These 8 regressions have a mean $R^2 =$<br>0.174 and mean $p = 0.326$ | | Not applicable for method<br>C, as species richness is<br>a property of a group, not<br>of species. |
| 2. Testing the hypothesised negative association between **phenotypic rates** and **branch duration**.<br>Expected given this relationship can be widely observed. **<u>Moderate support</u>.** |  |  |  |  |
| 75/125 (60%) of comparisons support a negative association<br>Negative association is particularly strong when comparing specific<br>comparisons, where 9/12 (75%) of Euryhaline vs. Freshwater-brackish comparisons, and 14/19 (73.7%) of Marine-brackish vs.<br>Marine comparisons support a neg. association. | | 8/10 regressions support a negative association. These<br>8 regressions have a mean $R^2 = 0.07$, mean $p = 0.558$ | | branch length $\log_{10} \sim$<br>size rate<br>($S = -0.361$, $p = 0$) |
| 3. Testing the hypothesised negative association between **phenotypic rates** and **size**.<br>Reasonable to hypothesise that, owing to the larger carrying capacity at lower trophic levels, smaller taxa may have a larger propensity for speciation than<br>larger taxa. **No support for a negative association from approaches A and B, yet a <u>significant positive association</u> with approach C.** |  |  |  |  |
| 63/125 (50.4%) support a neg. association. The negative association<br>is particularly poorly supported for Freshwater-brackish vs. Marine<br>comparisons, where support only occurs in 2/8 (25%) comparisons. | | 6/10 regressions support a negative association. These<br>6 regressions have a mean $R^2 = 0.043$, mean $p = 0.676$ | | Size $\log_{10} \sim$ beta $\log_{10}$<br>($S = 0.21$, $p = 0.002$) |

Approach A (Table 1A) determines if each theorised association in Table 1 is plausibly active (yes/no) in each of the 125 comparisons (Appendix 5). For example, to test for support for the hypothesised negative association between rates of size evolution and branch duration in just one comparison (i.e. the comparison between marine and freshwater Gobies), we ask whether the habitat with the shorter mean tip-branch durations also possesses the higher mean rate of size evolution. If true, this single comparison supports the negative branch duration vs. rate association. We then ask what percentage of results support the association, and because these 125 comparisons are a mixture of all 10 sets of pairwise habitat comparisons (e.g. all the marine vs. freshwater represent one set comparing two habitats), we also obtained percentage support for each set (Appendix 6) to reveal if specific associations are prevalent for specific environmental comparisons (e.g. if shorter branch durations only correlate with higher size rates in marine vs. freshwater comparisons). Where percentages for any habitat pairs were notably high/low, these are presented in Table 1A).

Approach B tested the three associations in a continuous manner, to determine the extent to which increases in one variable (when comparing between two habitats) are met by corresponding increases/decreases in the second variable; this determines the strength of association. For each variable, a regression was performed for each of the 10 pairwise habitat comparisons (i.e. all order-level comparisons between marine and freshwater habitats comprise one regression) to determine if the relationship changes depending upon the habitats compared (Appendices 7–9, summarised in Appendix 10). Table 1B displays information on whether a majority of the 10 regressions were positive or negative, and the average R^2^ and *p* values for whichever sign formed the majority (e.g. average of all positive associations). Approach C tests for an association between rates of size evolution and absolute body size with the full dataset using STRAPP (only applicable to species/tip variables).

Finally, we tested whether lacustrine species exhibited higher rates of phenotypic change (statistical results in Appendix 11) using the scheme applied by Miller (2021) which discovered higher speciation rates in lakes.

## RESULTS

### Rates of size evolution between salinity habitats

11k-tree full-dataset distributions of log size rates show minor differences between habitats (Fig. 1A), with only the median rates for freshwater-brackish taxa appearing discernibly higher that median tip rates for other salinities. Of the four phylogenetic methods applied that are capable of performing significance tests, none of them, applied to either the untransformed (Appendix 1) or logged (Appendix 2) full dataset rate data, recover significant differences between any two habitats. With these methods, only one significant difference emerges between any habitats with the full 31k-tree dataset (Fig. 1B): higher rates in freshwater-brackish taxa vs. marine-brackish taxa (*p* = 0.007, compare.rates; Appendix 3).

**FIGURE 1.**
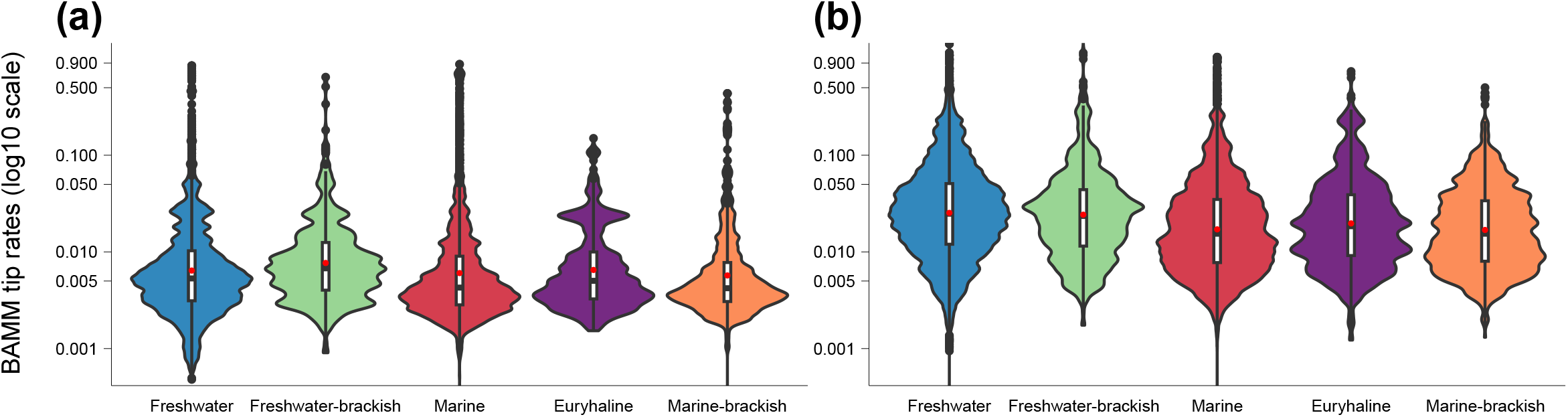
Distributions of phenotypic log10 tip rates for taxa in each salinity habitat using (a) the 11k-tree dataset and (b) the 31k-tree dataset. Tip rates represent the Brownian rate parameter with units of log(cm)^2 / myr.

Phenotypic rates in the 31k-tree dataset are inflated relative to the 11k-tree dataset in every habitat (Fig. 1), which may arise due to phylogenetic structure in the additional taxa in the supertree, and also, potentially, due to the smaller sizes of those additional taxa, as reported in Clarke (2021). Elevation of freshwater species rates is sufficient for their median rate to match that of freshwater-brackish habitats.

Rate differences between habitats were assessed for each unique pair of habitat comparisons (e.g. marine vs. freshwater) by i. calculating the percentage of clades where mean rate in one habitat is larger than the other at each phylogenetic scale (e.g. for families containing both marine and freshwater species, asking in what percentage of these families the marine members possess higher rates); and ii. whether patterns regarding these percentages are repeated at multiple phylogenetic scales. Thus, if there were a highly consistent pattern of rate difference between any two habitats (i.e. where most clades at most scales tended to shower higher rates in one habitat relative to another), we expect individual rows of Fig. 2A to be dominated by a single colour. However, rows are rarely dominated by a single colour, illustrating a lack of consistency regarding which habitats possess higher rates than others. Thus, rate patterns vary considerably with clade and scale. Only one finding – higher rates in freshwater-brackish taxa vs. euryhaline taxa – consistently emerges in a majority of examined clades at most scales (i.e. in over 50% of comparisons within each scale, but most commonly for 70–100% of comparisons); this is the case for all but one phylogenetic scale using both the 11k- and 31k-tree datasets (Fig. 2A–C, row 3). This finding is maintained regardless of which mean rate metric (untransformed tip-rate, log10 tip-rate, PGLS tip-rate and compare.rates means) is used to compares rates of size evolution (see row 3 in the following figures: Fig. 2, pages 2, 4, 10 and 12 of Appendix 1, page 2 of Appendix 2). However, even in this strongest example, this finding does not emerge for evolutionary hotspot comparisons, where either exactly 50% of the groupings examined show higher freshwater-brackish rates relative to euryhaline rates (Fig. 2A-B, row 3) or even less than 50% (compare.rates function, see row 3 of pages 2, 4, 10 and 13 of Appendix 1 and page 2 of Appendix 2). This shows the potential for “evolutionary hotspot” clades (i.e. that explore many salinities) to show atypical patterns relative to non-hotspot clades, as has been seen for body size patterns (Clarke, 2021). Tests for statistical significance are made with several metrics (Appendices 1–4). The second strongest finding is that within many (but not all) phylogenetic scales, there is a small majority of clades (∼55–65%) for which freshwater taxa possess higher mean rates than marine taxa – a pattern that can strengthen with some 31k-tree metrics (see row 8 in the following figures: Fig. 2, pages 2, 4, 10 and 13 of Appendix 3, page 2 of Appendix 4).

**FIGURE 2.**
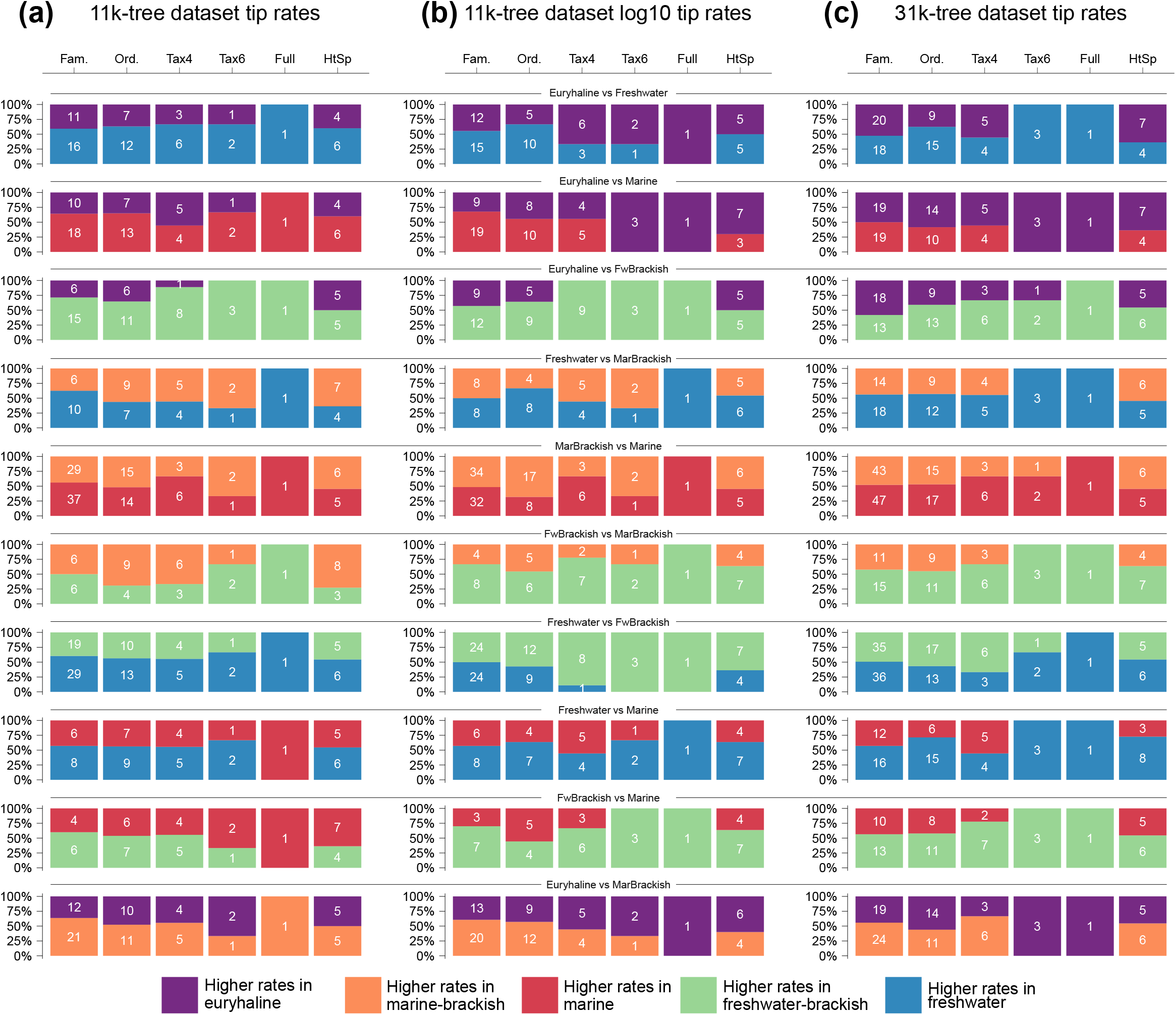
The percentage of clades where the average phenotypic rate in one habitat is larger than a second habitat to which it is being compared. Each row depicts comparison of two habitats, and white numbers capture how many groupings supported higher rates for a specific habitat. For example, row 1 column 1 in (a) shows that of the 27 fish families containing both euryhaline and freshwater species, the freshwater species showed higher rates of body size evolution in 59% of these families (i.e. 16 of 27). Comparisons are made between every unique pair of habitats (from five habitat types) across 10 scales of observation (6 shown, Appendices 1–4 show results for all 10 scales using all examined rate metrics). The panels correspond to comparisons of (a) phenotypic rate upon the 11k-tree dataset; (b) log10 phenotypic rate upon the 11k-tree dataset; and (c) log10 phenotypic rate upon the 31k-tree dataset. Presentation of logged and untransformed rate data shows how this decision impacts inferred patterns; firm conclusions are only drawn when all metrics agree. Taxonomic scale definitions provided in methods. HtSp = evolutionary hotspots.

### Alternative drivers of size rates

Table 1 summarises the degree of support recovered for three theorised rate drivers across three approaches. Together, these inform a relative ranking, from the strongest (1) to the weakest (3) as listed in Table 1 alongside a support statement. Thus, phenotypic rates are best correlated with species richness, followed by branch duration, then absolute size. As an illustrative example, we here interpret the results for the association ranked 1^st^ – the hypothesised positive association between phenotypic rates and species richness (Table 1).

Method A shows positive associations occur in a minority of comparisons. Specifically, in 38.4% of the 125 order-level comparisons, the group with higher species richness also shows higher rates of phenotypic evolution (Appendices 5–6). The text in Table 1A, reports noteworthy results, showing this percentage can fall as low as 20% when examining the 12 freshwater-brackish vs. marine-brackish comparisons or the 18 euryhaline vs. marine-brackish comparisons. Beyond reading Appendices 5–6, one can visually assess which habitat comparisons at the order level show a positive or negative association between rates and species richness in Appendix 7, where each of the 10 plots depicts contrasts in richness and rates between two habitats; points in grey quadrants support a negative association while points in white quadrants support a positive association. These 10 contrast plots form the basis of the 10 regressions that define method B. As for method A, the hypothesised positive association is rejected, as only 2 of the 10 regressions find any positive relationship (with a tiny average R^2^ value of 0.02), while 8 of 10 show a negative association with an average R^2^ of 0.17. Thus, while the hypothesised positive association is rejected, the opposite, negative association is supported. The average R^2^ value for negative associations is larger than for any other variable tested, thus this variable is ranked 1^st^. Method C, which involves a full dataset STRAPP analysis can only be applied to species-level traits, and therefore cannot be applied to species richness. Figure 3 illustrates the negative and positive relationships between tip rates with tip-branch duration and body size, respectively, to which STRAPP analyses could be applied (Table 1C).

**FIGURE 3.**
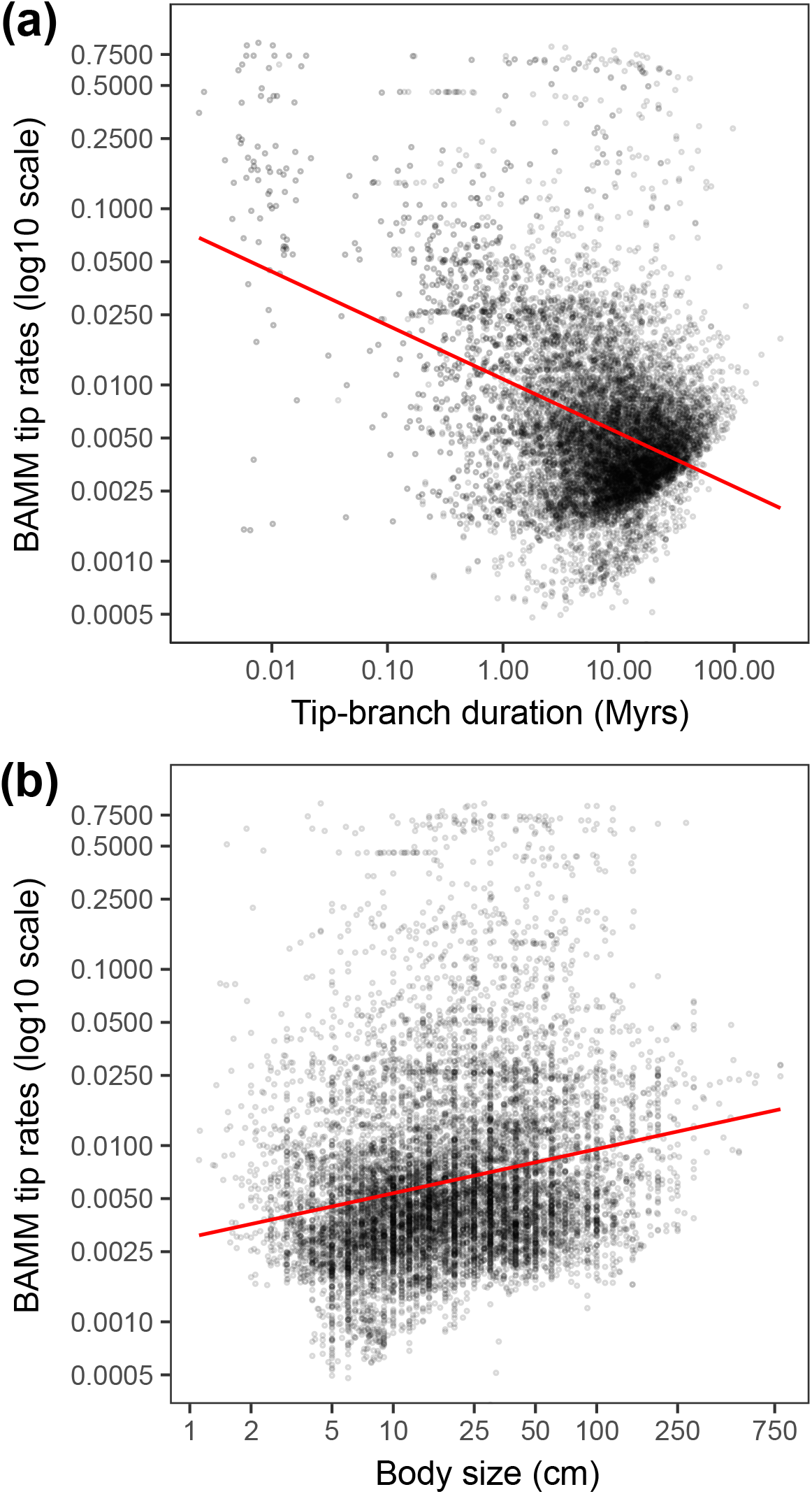
Scatter plots of BAMM tip rates for body size evolution on a log10 scale as a response variable to (a) tip-branch duration and (b) body size. Regression lines are for illustrative purposes, more appropriate statistical outputs derived from STRAPP are reported in Table 1C. Tip rates represent the Brownian rate parameter with units of log(cm)^2/ myr.

Finally, we assessed the role of an alternative habitat scheme upon rates of phenotypic evolution to check if lacustrine environments elevate rates of phenotypic evolution as they do for speciation rates (Miller, 2021). We confirm this to be true; exclusively lacustrine taxa possess the highest mean and median rates, followed by simultaneously lacustrine + riverine taxa, followed by taxa in three habitats with comparable averages (diadromous taxa that migrate between saltwater and freshwater, exclusively marine, exclusively riverine; Fig. 4, Appendix 11). Lacustrine taxa possess statistically higher rates than marine and riverine taxa (STRAPP; Appendix 11). Furthermore, simultaneously lacustrine + riverine taxa possess statistically higher rates than exclusively riverine taxa (log10 rates phylogenetic ANOVA and STRAPP; Appendix 11).

**FIGURE 4.**
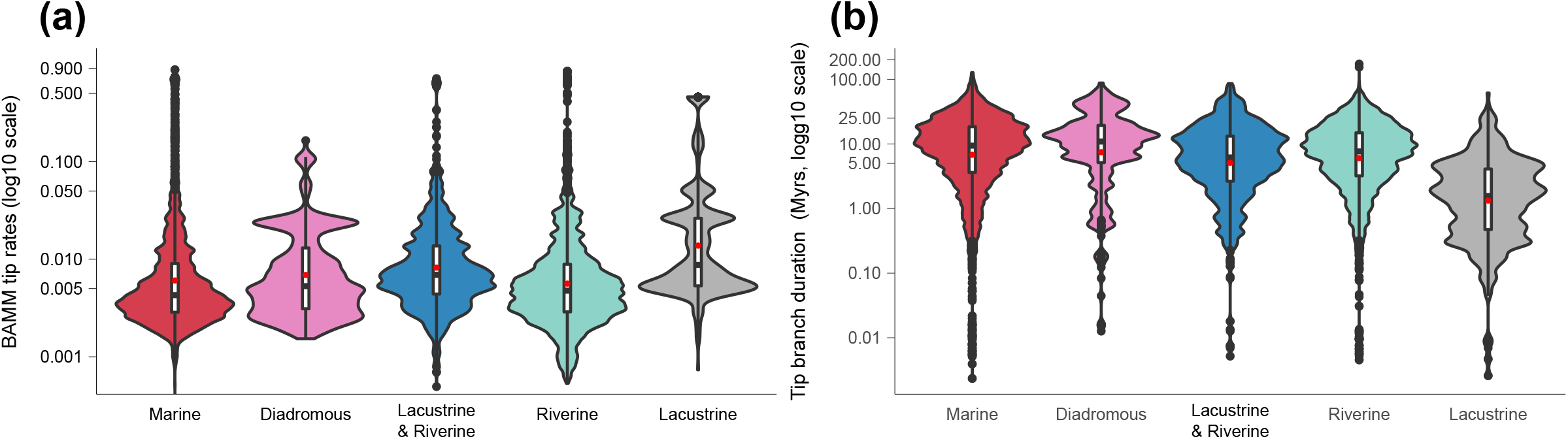
Distributions of (a) phenotypic log10 tip rates, and (b) log10 tip-branch durations, for taxa in the alternative habitat categories of Miller 2021. Tip rates represent the Brownian rate parameter with units of log(cm)^2 / myr.

## DISCUSSION

### Mostly limited role for salinity habitat in driving rate differences

The effects of salinity upon rates of body size evolution have been explored previously in different clades and at different scales, with variable statistical methodologies and habitat schemes, most commonly with a binary setup comparing marine and freshwater. Results have varied, showing: i. higher rates in freshwater clades, including pufferfishes (Tetraodontiformes, Santini *et al*., 2013), grunters (Terapontidae, Davis *et al*., 2014), and ‘goby’ clades (Apogonidae and Gobiidae + Gobionellidae, Thacker, 2014); ii. higher rates in marine clades, including sea catfishes (Ariidae; Betancur-R. *et al*., 2012) and Upper Jurassic–end Palaeocene actinopterygian fishes (Guinot and Cavin, 2018); and iii. no clear evidence for rate differences in needlefishes (Belonidae, Kolmann *et al*., 2020). Our experimental design aimed to reconcile these previous findings by repeatedly testing for differences in size rates between six salinity habitats at ten phylogenetic scales, allowing the reader to compare our results (using a variety of metrics) to most results found in previous studies (e.g. for any family or order of fishes (e.g. our Tetraodontidae, Belonidae and terapontid results in Appendices 1–4).

Despite classic literature characterising freshwaters as slow-evolving environments (Darwin, 1877, 1859; Darwin and Seward, 1903), and several contemporary efforts highlighting freshwaters as dynamic, high-paced environments for evolution (May and Godfrey, 1994; Grosberg *et al*. 2012; Tedesco *et al*., 2017), neither marine nor freshwater taxa show higher mean rates in a majority of clades at any of the phylogenetic scales examined (row 8 in both Fig. 2 and Appendices 1–4). Even for the whole dataset, whether freshwater or marine taxa possess the higher rate varies with dataset and metric. Nevertheless, freshwater taxa do have a mild rate advantage over marine taxa, showing higher rates in small majorities (∼55–65%) of clades when the 11k-tree data is examined at many taxonomic scales (e.g. Family, Order, Tax6), making it the second most consistent pattern recovered in the study; this pattern becomes stronger on 31k trees, despite the fact that marine and freshwater taxa are presented by comparable proportions of species in both the 11k molecular trees and 31k supertrees, suggesting that any supertree bias towards higher rates cannot fully explain this freshwater rate advantage. However, exceedingly few individual comparisons at any scale are statistically significant when phylogenetic tests are applied (Appendices 1–4).

The inclusion of several less explored salinity habitats (e.g. euryhaline, freshwater-brackish, marine-brackish), mostly did little to reveal strong and consistent rate differences between any pair of habitats. Only one full dataset comparison recovered a significant rate difference with one dataset and phylogenetic method (higher rates in freshwater-brackish taxa verses marine in the 31k-tree dataset with compare.rates; Appendices 3–4). In nine of ten habitat comparisons, no consistent pattern of differences between two rates occurred in a majority of clades within most phylogenetic scales, suggesting that, unlike patterns of absolute body size between these same salinity settings (which are consistent and clear across scales and dataset types, (Clarke, 2021), rate difference patterns are mostly clade- and scale-dependent. Size rate patterns therefore echo two claims made for speciation rates (Miller, 2021; Rabosky, 2020) – firstly, differences between marine and freshwater environments are relatively subtle, and by comparison, size rates are even more subtle given that higher speciation in freshwater taxa can achieve statistical significance; secondly, results seen at the full dataset level are often driven by specific clades (higher speciation rates in freshwater fishes driven by Cichliformes), which occurs in our data frequently, such as when any size rate pattern at a higher scale does not match the result seen in over 50% of comparisons at a lower scale.

One novel difference, which emerged in a majority of clades analysed for most taxonomic scales (with all four rate metrics and in the 31k-tree dataset), is higher rates in freshwater-brackish taxa relative to their euryhaline relatives (row 3 in Fig 2). Higher phenotypic rates may, in part, relate to the fact that freshwater-brackish taxa comprise large numbers of species from opposite ends of the body-size distribution. Clarke (2021) showed that freshwater-brackish settings receive ∼362 transitions from freshwater (where species are the smallest relative to other salinity habitats) and a combined ∼175 transitions from euryhaline and marine-brackish habitats, which are jointly the largest on average; this can be compared to freshwater-brackish settings, which show medium sizes on average. As such, higher rates may occur as arrivals from habitats with species at opposite ends of the size spectrum evolve towards medium body sizes, consistent with observations that larger amounts of size change occur when fishes move between (rather than within) salinity habitats, a pattern that occurs particularly when transitioning into brackish/mixed settings (Guinot and Cavin, 2018). Furthermore, given that the number of transitions into freshwater-brackish is high relative to the total species richness of the habitat and also exceeds the number of outward transitions (∼557 in, ∼369 out), it is plausible that lineages in neighbouring habitats that are the most likely to transition into freshwater-brackish are those already experiencing elevated speciation rates and rates of body size evolution, and thus, bring those high rates with them, increasing the average rate of freshwater-brackish lineages. By contrast, the euryhaline lifestyle (compared to which freshwater-brackish taxa consistently show higher rates) loses more lineages than it gains via such transitions (∼535 in, ∼700 out), and so may frequently lose lineages exhibiting high rates of size evolution to other environments.

From the other variables examined here, it is plausible to ask whether the lower species richness (Appendix 7), shorter branch durations (Fig. 3A, Appendix 8), or smaller body sizes (Fig. 3B, Appendix 9) of freshwater-brackish species can explain their higher rates relative to euryhaline one. While shorter branch durations and lower species richness may theoretically play a role (given their broader significance in driving rates, detailed below), it is unlikely that smaller size of freshwater-brackish taxa relative to euryhaline taxa is responsible for higher rates in freshwater-brackish taxa, as phenotypic rates tend to gently increase as body size increases (detailed below). Shorter branch durations may play a small role, as 3/8 of the order-level comparisons where freshwater-brackish taxa report higher rates than euryhaline relatives also possess shorter mean tip-branch durations (e.g. for Cyprinodontiformes and Siluriformes; Appendix 8). Lower species richness has more potential, as 6/8 of the higher-rate freshwater-brackish orders also possess lower species richness (yet is not true for Cyprinodontiformes and Siluriformes; Appendix 7). Thus, it is possible that both these factors act alongside high inwards transition rates and evolution from extremes towards medium sizes (both discussed above) to help deliver higher rates in freshwater-brackish taxa.

### Determining the relative importance of additional drivers of size rate variation

Given the limited ability for salinity to predict differences in size rates, we evaluated alternative drivers with three methods (Table 1). One can check whether each driver is plausibly active for all 1056 habitat and clade comparisons made with the 11k-tree dataset (Appendix 5). Taken together, the best supported association is the negative relationship between species richness and size rates. We hypothesised a positive relationship based upon the known association between phenotypic rates and speciation rates (Rabosky et al., 2013) combined with an expectation that more species-rich groups (especially in the confines of our within-order comparisons, where smaller differences in branch duration can accrue) should generally be underpinned by higher speciation rates. Yet support for a positive association between size rates and species richness was not compelling, seen in just ∼38% of comparisons and only 2 of the 10 regression analyses (Table 1A–B). Within our 125 order-level comparisons, it is rare for any habitat to display both high mean species richness and mean size rates simultaneously relative to a comparator group (i.e. if they existed, they would plot towards the outer corners of white quadrants in Appendix 7). Nevertheless, 7 comparisons notably combine moderately high size rates with moderately high species richness, including marine Perciformes (vs. freshwater, marine-brackish, and euryhaline relatives), freshwater-brackish Cyprinodontiformes (vs. euryhaline), euryhaline Clupeiformes (vs. freshwater-brackish), freshwater Siluriformes (vs. marine-brackish) and freshwater Characiformes (vs. freshwater-brackish). These 7 outcomes may indicate where diversity-dependent mechanisms (i.e. where speciation rate decreases as the number of species increases; e.g. Rabosky, 2013; Rineau *et al*. 2022) are either absent or simply do not impede higher phenotypic rates, highlighting them as good candidates for further study on this question.

Beyond these 7 exemplars, most comparisons (∼62%) show that the habitat with the higher species richness possesses lower mean size rates across, and many display a substantial degree of mismatch, explaining why 8/10 regressions report negative associations (Table 1B, Appendix 7). The absolute largest mismatch observed (i.e. combining high rates with low richness) is seen in marine-brackish Cyprinodontiformes (vs. freshwater), followed by a host of substantial mismatches, including freshwater Tetraodontiformes (vs. marine), marine-brackish Cyprinodontiformes (vs. freshwater-brackish), marine-brackish Labriformes (vs. marine), euryhaline Chaetodontiformes (vs. marine-brackish), freshwater-brackish Perciformes (vs. freshwater) and numerous others (see points falling within central regions of grey quadrants in Appendix 7). In all comparisons showing mismatch, it is therefore plausible that diversity-dependent factors act to supress phenotypic rates and/or ecological opportunity in the more species-rich groupings, and therefore we flag these groups for future study. Beyond these strong exemplars, the high prevalence of negative associations suggests these processes may be commonplace in fishes.

The second strongest association is the negative relationship between size rates with branch duration (Table 1). The negative time scaling of evolutionary rates – where observed rates decrease as the time periods over which they are measured increase – appears to be a pervasive feature in evolutionary biology, with explanations ranging from incorrect model assumptions or artifacts as a result of estimation errors and/or sampling bias, to acceptance that the pattern is real (Harmon et al., 2021). Our study therefore confirms the presence of this phenomenon in the half of vertebrate diversity represented by fishes (Fig. 3A). However, substantial rate variation occurs around the proportion explained by branch duration. Thus, while it is generally true that a very long branch will possess a lower rate than a very short branch, if the branches being compared are not orders of magnitude different in duration, the shorter branch might not show higher rates than the longer one. This explains why STRAPP (which considers the whole dataset) shows a strong, significant negative relationship, while this negative association is weak with approaches A and B (where 60% of binary comparisons show negative associations, and the negative regressions average a small R^2^ of 0.07, respectively; Table 1A–B), because approaches A and B compare habitats within orders that generally do not differ by large amounts in their average tip-branch durations. When moderate to large differences in tip-branch durations do emerge (i.e. points that plot further away from 0 on the x axis in Appendix 8), most support a negative association (i.e. they plot within grey quadrants) which produces negative relationship regressions. Notably, the shorter tip-branch durations of lacustrine taxa habitats appear to underpin their higher phenotypic rates relative to other habitats (Fig 4). In notably rare cases, habitats show both substantially longer average tip-branch durations and higher rates than their relatives from another habitat, including marine-brackish Cyprinodontiformes (vs. euryhaline) and euryhaline Chaetodontiformes (vs. marine) (Appendix 8).

Although we had predicted that smaller taxa would possess higher rates of size evolution, we instead found a significant positive relationship between body size and rate using STRAPP on the full dataset (Table 1C, S=0.21, *p*=0.002). However, results were evenly split between positive and negative associations with approaches A and B (Table 1A–B). The failure of approaches A and B to clearly support a positive relationship can be explained by the low Spearman’s value (0.21) in approach C, revealing that substantial variation in rate is not explained by size (Fig 3B). As such, the differences in mean size between any comparisons in approaches in A and B would regularly need to be very large to frequently detect a positive association, which is unlikely for within-order comparisons.

A positive association between size and size rates, where rates of size evolution increase as absolute body size increases, agrees with equivalent findings in both mammals (Monroe and Bokma, 2009) and perches (Arbour and Stanchak, 2021). Yet, closer examination reveals key features. First, the very highest rates occur in the middle of the size distribution (Fig. 3B), although this may occur because most species reside in this region, and so there are more opportunities for higher rates to arise by chance. Second, the distribution of rates appears to shift suddenly to higher rates in both the very largest and very smallest taxa (Fig. 3B). Specifically, the very lowest rates occur in fishes between 5 and 10 cm long, but below 5 cm, rates increase suddenly forming a distribution of low-medium rates, and then, at sizes below 2 cm, to rates close to 0.01 and above. Similarly, at the very largest sizes, only rates of around ∼0.0025 occur. Finding higher size rates at body size extremes matches the higher size rates found at trophic extremes in reef fishes (Borstein et al., 2019). Thus, several nuanced relationships can occur between size and size rates even within the context of a broad positive association.

Finally, we considered the impact of lakes on phenotypic rates by implementing the alternative habitat scheme Miller (2021), and found significantly higher rates in lacustrine habitats relative to marine and riverine taxa (Appendix 11), confirming that patterns seen for speciation rate (Miller, 2021) are matched by size rates. Importantly, elevated rates in lakes are not an isolated phenomenon restricted to cichlids, but is observed in a wide variety of orders (Appendix 12), most notably for Cyprinodontiformes, Clupeiformes, Siluriformes, Cypriniformes, Osmeriformes, Beloniformes, and Atheriniformes, though it is absent for Osteoglossiformes, Perciformes, and Synbranchiformes. Elevated rates in lakes are consistent with evolutionary theory, as lake radiations are commonly cited as providing ecological opportunity for adaptive radiations of species, characterised by rapid speciation and phenotypic differentiation (Schluter, 2000; Simpson, 1953). Furthermore, we expect phenotypic rates to be enhanced by both the greater depth profile of lakes, presenting greater habitat variety (Lucek et al., 2016; Recknagel et al., 2017), and the shorter tip-branch durations of lacustrine species (Fig. 4B), as well as the potential for phytoplankton-based food webs in lacustrine systems, which can promote longer food chains (Potapov et al., 2019), promoting greater fish body size variety (Clarke, 2021).

In conclusion, rates of body size evolution rarely show consistent differences between salinity habitats in a way that is maintained for most clades examined at most phylogenetic scales. By examining many clades, we can reveal that disagreement between previous studies is in fact a genuine reflection of the highly variable nature of the pattern, where the number of clades for one outcome are counterbalanced by a similar number of clades showing the opposite outcome. However, freshwater-brackish taxa do possess consistently higher rates compared with euryhaline taxa at all scales examined. Tests of additional drivers of rate variation reveal that low species richness and short tip-branch durations are associated with higher rates of size evolution, with more subtle evidence of higher rates associated with greater absolute body size. Finally, we found significantly elevated rates of size evolution within lacustrine species relative to all other habitats, and species straddling both rivers and lakes relative to exclusively riverine species. Together, our analyses reveal the rate patterns that underpin global patterns of body size for fishes, identifying both factors that play a limited role (i.e. salinity) and those that explain considerable amounts of rate variation across half of vertebrate diversity.

## REFERENCES

Adams, D.C., Otárola-Castillo, E., 2013. geomorph: an r package for the collection and analysis of geometric morphometric shape data. Methods Ecol. Evol. 4, 393–399. 10.1111/2041-210X.12035

Arbour, J.H., Stanchak, K.E., 2021. The little fishes that could: smaller fishes demonstrate slow body size evolution but faster speciation in the family Percidae. Biol. J. Linn. Soc. 134, 851–866. 10.1093/biolinnean/blab125

Betancur-R., R., Ortí, G., Stein, A.M., Marceniuk, A.P., Alexander Pyron, R., 2012. Apparent signal of competition limiting diversification after ecological transitions from marine to freshwater habitats. Ecol. Lett. 15, 822–830. 10.1111/j.1461-0248.2012.01802.x

Blackburn, T.M., Gaston, K.J., 2002. Scale in macroecology. Glob. Ecol. Biogeogr. 11, 185–189. 10.1046/j.1466-822X.2002.00290.x

Borstein, S.R., Fordyce, J.A., O’Meara, B.C., Wainwright, P.C., McGee, M.D., 2019. Reef fish functional traits evolve fastest at trophic extremes. Nat. Ecol. Evol. 3, 191–199. 10.1038/s41559-018-0725-x

Clarke, J.T., 2021. Evidence for general size-by-habitat rules in actinopterygian fishes across nine scales of observation. Ecol. Lett. 24, 1569–1581. 10.1111/ele.13768

Collyer, M.L., Adams, D.C., 2018. RRPP: An r package for fitting linear models to high-dimensional data using residual randomization. Methods Ecol. Evol. 9, 1772–1779. 10.1111/2041-210X.13029

Cooney, C.R., Thomas, G.H., 2020. Heterogeneous relationships between rates of speciation and body size evolution across vertebrate clades. Nat. Ecol. Evol. 5, 101–110. 10.1038/s41559-020-01321-y

Darwin, C., 1877. Descent of man, and selection in relation to sex. Revised and augmented 2nd edition, including an essay by T. H. Huxley. John Murray, London.

Darwin, C., 1859. On the origin of species by means of natural selection, or, The preservation of favoured races in the struggle for life. John Murray, London.

Darwin, F., Seward, A.C., 1903. More Letters of Charles Darwin: a record of his work in a series of hitherto unpublished letters. London.

Davis, A.M., Unmack, P.J., Pusey, B.J., Pearson, R.G., Morgan, D.L., 2014. Effects of an adaptive zone shift on morphological and ecological diversification in terapontid fishes. Evol. Ecol. 28, 205–227. 10.1007/s10682-013-9671-x

Finlay, B.J., Thomas, J.A., McGavin, G.C., Fenchel, T., Clarke, R.T., 2006. Self-similar patterns of nature: insect diversity at local to global scales. Proc. R. Soc. B Biol. Sci. 273, 1935–1941. 10.1098/rspb.2006.3525

Foody, G.M., 2004. Spatial nonstationarity and scale-dependency in the relationship between species richness and environmental determinants for the sub-Saharan endemic avifauna. Glob. Ecol. Biogeogr. 13, 315–320. 10.1111/j.1466-822X.2004.00097.x

Froese, R., Pauly, D. (eds), 2000. FishBase 2000: Concepts, designs and data sources. WorldFish.

Grosberg, R.K., Vermeij, G.J., Wainwright, P.C., 2012. Biodiversity in water and on land. Curr. Biol. 22, R900–R903. 10.1016/j.cub.2012.09.050

Guinot, G., Cavin, L., 2018. Body size evolution and habitat colonization across 100 million years (Late Jurassic– Paleocene) of the actinopterygian evolutionary history. Fish Fish. 19, 577–597. 10.1111/faf.12275

Harmon, L.J., Pennell, M.W., Henao-Diaz, L.F., Rolland, J., Sipley, B.N., Uyeda, J.C., 2021. Causes and Consequences of Apparent Timescaling Across All Estimated Evolutionary Rates. Annu. Rev. Ecol. Evol. Syst. 52, 587–609. 10.1146/annurev-ecolsys-011921-023644

Igea, J., Miller, E.F., Papadopulos, A.S.T., Tanentzap, A.J., 2017. Seed size and its rate of evolution correlate with species diversification across angiosperms. PLOS Biol. 15, e2002792. 10.1371/journal.pbio.2002792

Kolmann, M.A., Burns, M.D., Ng, J.Y.K., Lovejoy, N.R., Bloom, D.D., 2020. Habitat transitions alter the adaptive landscape and shape phenotypic evolution in needlefishes (Belonidae). Ecol. Evol. 10, 3769–3783. 10.1002/ece3.6172

Lee, C.E., Bell, M.A., 1999. Causes and consequences of recent freshwater invasions by saltwater animals. Trends Ecol. Evol. 14, 284–288. 10.1016/S0169-5347(99)01596-7

Lucek, K., Kristjánsson, B.K., Skúlason, S., Seehausen, O., 2016. Ecosystem size matters: the dimensionality of intralacustrine diversification in Icelandic stickleback is predicted by lake size. Ecol. Evol. 6, 5256–5272. 10.1002/ece3.2239

May, R.M., Godfrey, J., 1994. Biological Diversity: Differences between Land and Sea [and Discussion]. Philos. Trans. Biol. Sci. 343, 105–111.

McClain, C.R., Nekola, J.C., 2008. The role of local-scale processes on terrestrial and deep-sea gastropod body size distributions across multiple scales. Evol. Ecol. Res. 10, 129–146.

Miller, E.C., 2021. Comparing diversification rates in lakes, rivers, and the sea. Evolution 75, 2055–2073. 10.1111/evo.14295

Monroe, M.J., Bokma, F., 2009. Do Speciation Rates Drive Rates of Body Size Evolution in Mammals? Am. Nat. 174, 912–918. 10.1086/646606

Potapov, A.M., Brose, U., Scheu, S., Tiunov, A.V., 2019. Trophic Position of Consumers and Size Structure of Food Webs across Aquatic and Terrestrial Ecosystems. Am. Nat. 194, 823–839. 10.1086/705811

Rabosky, D.L., 2020. Speciation rate and the diversity of fishes in freshwaters and the oceans. J. Biogeogr. 47, 1207–1217. 10.1111/jbi.13839

Rabosky, D.L., 2015. No substitute for real data: A cautionary note on the use of phylogenies from birth–death polytomy resolvers for downstream comparative analyses. Evolution 69, 3207–3216. 10.1111/evo.12817

Rabosky, D.L., 2013. Diversity-Dependence, Ecological Speciation, and the Role of Competition in Macroevolution. Annu. Rev. Ecol. Evol. Syst. 44, 481–502. 10.1146/annurev-ecolsys-110512-135800

Rabosky, D.L., Chang, J., Title, P.O., Cowman, P.F., Sallan, L., Friedman, M., Kaschner, K., Garilao, C., Near, T.J., Coll, M., Alfaro, M.E., 2018. An inverse latitudinal gradient in speciation rate for marine fishes. Nature 559, 392–395. 10.1038/s41586-018-0273-1

Rabosky, D.L., Grundler, M., Anderson, C., Title, P., Shi, J.J., Brown, J.W., Huang, H., Larson, J.G., 2014. BAMMtools: an R package for the analysis of evolutionary dynamics on phylogenetic trees. Methods Ecol. Evol. 5, 701–707. 10.1111/2041-210X.12199

Rabosky, D.L., Huang, H., 2016. A Robust Semi-Parametric Test for Detecting Trait-Dependent Diversification. Syst. Biol. 65, 181–193. 10.1093/sysbio/syv066

Rabosky, D.L., Santini, F., Eastman, J., Smith, S.A., Sidlauskas, B., Chang, J., Alfaro, M.E., 2013. Rates of speciation and morphological evolution are correlated across the largest vertebrate radiation. Nat. Commun. 4, 1958. 10.1038/ncomms2958

Rahbek, C., 2005. The role of spatial scale and the perception of large-scale species-richness patterns. Ecol. Lett. 8, 224–239. 10.1111/j.1461-0248.2004.00701.x

Recknagel, H., Hooker, O.E., Adams, C.E., Elmer, K.R., 2017. Ecosystem size predicts eco-morphological variability in a postglacial diversification. Ecol. Evol. 7, 5560–5570. 10.1002/ece3.3013

Revell, L.J., 2012. phytools: an R package for phylogenetic comparative biology (and other things). Methods Ecol. Evol. 3, 217–223. 10.1111/j.2041-210X.2011.00169.x

Rineau, V., Smyčka, J., Storch, D., 2022. Diversity dependence is a ubiquitous phenomenon across Phanerozoic oceans. Sci. Adv. 8, eadd9620. 10.1126/sciadv.add9620

Santini, F., Nguyen, M.T.T., Sorenson, L., Waltzek, T.B., Lynch Alfaro, J.W., Eastman, J.M., Alfaro, M.E., 2013. Do habitat shifts drive diversification in teleost fishes? An example from the pufferfishes (Tetraodontidae). J. Evol. Biol. 26, 1003–1018. 10.1111/jeb.12112

Schluter, D., 2000. Ecological Character Displacement in Adaptive Radiation. Am. Nat. 156, S4–S16. 10.1086/303412

Simpson, G.G., 1953. The Major Features of Evolution. Columbia University Press, New York Chichester, West Sussex. 10.7312/simp93764

Slater, G.J., Friscia, A.R., 2019. Hierarchy in adaptive radiation: A case study using the Carnivora (Mammalia). Evolution 73, 524–539. 10.1111/evo.13689

Tedesco, P.A., Paradis, E., Lévêque, C., Hugueny, B., 2017. Explaining global-scale diversification patterns in actinopterygian fishes. J. Biogeogr. 44, 773–783. 10.1111/jbi.12905

Thacker, C.E., 2014. Species and shape diversification are inversely correlated among gobies and cardinalfishes (Teleostei: Gobiiformes). Org. Divers. Evol. 14, 419–436. 10.1007/s13127-014-0175-5

